# Exploring shaped focused ion beams for lamella preparation

**DOI:** 10.1101/2025.04.01.646235

**Authors:** Johann Brenner, Jürgen M. Plitzko, Sven Klumpe

**Affiliations:** Research Group CryoEM Technology, Max Planck Institute of Biochemistry, Martinsried, Germany; Research Institute of Molecular Pathology (IMP) and Institute of Molecular Biotechnology Austria (IMBA), Vienna BioCenter, Vienna, Austria

**Keywords:** Focused ion beams, beam shaping, lamella preparation, TEM, Cryo-FIB, Biology, cryo-electron tomography, patterning, ion knife

## Abstract

Focused ion beams (FIB) are widely used instruments in transmission electron microscopy (TEM) sample preparation across scientific disciplines. Generally, site-specific ablation of material is achieved by scanning a highly focused probe across a selected area, leading to the removal of material. However, the geometries of TEM lamellae milled with the FIB are usually highly non-isometric, with their thickness generally being orders of magnitude smaller than their width and length. Here, we explore a changed probe shape for milling. Instead of using a Gaussian-like spot probe ion beam, we characterize the use of the stigmator as quasi-cylindrical lens to create a highly astigmatic beam that we term ‘ion knife’. Using the ion knife allows for material ablation by spreading the current over a larger area and changes the dimension of the probe with an anisotropic change in apparent beam resolution, as observed in spot burn cross-sections. We demonstrate a method to approximate beam shapes by imaging that allows for convenient alignment of parameters for beam shaping. Finally, exploring shaped probes in cryogenic lamella preparation, we demonstrate the feasibility of cellular lamella milling and sectioning of cryo-lift-out volumes with the ion knife.

## 1. Introduction

Focused ion beams are widely used instruments in sample preparation of transmission electron microscopy (TEM) for the ultrastructural investigation of specimens ranging from materials [1] and semiconductors [2] to cells and organisms [3-6]. In these applications, the focused ion beam (FIB) is used as an imaging and site-specific shaping tool. The ions are usually generated from liquid metal ion sources (LMIS, commonly gallium) [7, 8] or plasma ion sources, mostly inductively coupled or electron cyclotron resonance plasma (ICP and ECR, respectively) from gases such as xenon, argon, nitrogen, and oxygen [9-11]. Numerous other ion sources such as liquid metal alloy ion sources (LMAIS) using e.g. GaBiLi [12], gas field ion sources (GFIS) [13] and many others have been or are being explored [14].

Though many other applications exist, reviewed elsewhere [14], a main use case for focused ion beams in sample preparation is the generation of thin sections referred to as lamellae by ablating material using FIBs that can be transferred into the TEM for imaging. Generally, these thin sections are not isometric in their geometry: while their thickness ranges from several tens of nanometers in the materials science [15] to hundreds of nanometers in the life sciences [16], their width and length is usually in the range of micrometers for both applications. In lamella preparation for cryo-electron tomography (cryo-ET), the typical lateral dimensions reach tens of micrometers. The site-specific ablation of material is performed using a focused probe that is scanned across the sample [17]. The atoms of the material surface are sputtered away by the incident ions, ablating the sample surface, resulting in a lamella. The volume of the ablated material depends on multiple parameters of the incident beam: atomic number, voltage, current, the material-specific properties of the specimen, and the utilized scanning strategy [18]. Additionally, the damage or implantation that the incident ions cause to the material is also a function of multiple factors, most notably their acceleration voltage, impact angle, and current density [19]. While most of these parameters have been explored extensively, to the best of our knowledge, none of the existing strategies of material ablation explore alterations to the shape of the incident beam.

In fluorescence light microscopy, light sheet microscopy (LSM) has gained momentum as it allows for imaging of biological specimen at minimal dose at an approximate optical sectioning of a confocal laser scanning microscopy [20]. The light sheet used for illumination in LSM is created with a cylindrical lens, which has two different focal lengths orthogonal to each other [21, 22]. This configuration causes the lens to focus light strongly in one direction while the orthogonal direction experiences little to no focusing.

Here, we demonstrate that in analogy to LSM, the ion beam stigmator commonly available in modern FIB instruments can be used as an astigmatic lens in a first approximation to a cylindrical lens. The resulting optics form an ion beam that resembles the shape of a knife, an astigmatic FIB probe that we call ‘ion knife’. The dimension of these probes is defined by two parameters: the length of the knife and its diameter, i.e. the probe size in the orthogonal direction of its length. By using the stigmator excitation in one direction, the minimal diameter of the ion beam probe can be reduced at the expense of increased length. We show that the generated probe can be used in lamella preparation, distributing the current limited by the beam-defining aperture across the ion knife such that the non-isometric nature of the lamella is taken into account during the milling process. Additionally, we demonstrate that the shaped ion beam can be used to prepare lamellae from frozen-hydrated cells for cryo-ET, although it currently poses several remaining challenges in this mode of implementation. Nevertheless, ion knives could potentially pave the way for a reduction in material loss in the preparation of lamellae from frozen hydrated material by cryo-lift-out by increasing the retention of material in recently developed sequential sectioning lift-out protocols [23, 24].

## 2. Materials and Methods

### 2.1 Sample freezing

The yeast strain FWY003 [25] was inoculated from overnight cultures in YPD medium (1% yeast extract, 2% peptone, and 2% glucose) to an OD_600_ of 0.15 and grown at 30 °C to an OD_600_ of 0.8. For plunge freezing of cells on grids, copper grids (200 Mesh Cu SiO2 R1/4, Quantifoil) were plasma cleaned for 1 min from each side. 4 µl of cell suspension was applied onto grids and plunged into a liquid ethane-propane mixture using an EM GP2 (Leica Microsystems) at a blotting time of 2 s, humidity 100%, a temperature of 22 °C and automatic plunging after blotting. Plunged grids were clipped into FIB-AutoGrid cartridges and stored in liquid nitrogen for further use.

The *C. elegans* L1 larvae samples were prepared as described previously [24]. In brief, high pressure freezing planchettes (Type B) coated with cetyl palmitate are used in a ‘waffle’-type preparation [26] to freeze a suspension of L1 larvae in M9 medium supplemented with 20% Ficoll 70 kDa on a copper 75 hexagonal mesh grid coated with formvar. High-pressure freezing was performed using a Leica EM Ice (Leica Microsystems). Grids were recovered and assembled into AutoGrids for subsequent experiments.

### 2.2 Theoretical model and experimental procedure for the generation of shaped ion beams using the FIB stigmator

The beam profile was simulated for a simple electrostatic quadrupole (ESQ) in python. The ESQ was modeled after derivations described in detail elsewhere by Masek and Sherwood [27]. A detailed theoretical background is provided within the jupyter notebook, available on GitHub (See Data and Code availability). Briefly, a two-dimensional Gaussian-like distribution (RMS = 250 nm) was used to approximate the FIB beam, sampling 10 million gallium ions. From this distribution, the particle trajectory as well as its axial derivative was calculated with a vectorized ESQ model in NumPy for a quadrupole length of 10 cm and a quadrupole strength parameter K. Finally, the resulting beam intensities were plotted in matplotlib for the stigmatic beam (K = 0 m^-1^) and the strongly astigmatic beam (K = -20 m^-1^).

Experimental implementation of beam shaping was performed on an Aquilos 2 and a Scios FIB-SEM system (Thermo Fisher Scientific) using a piece of a silicon wafer (WSM40525250P1324XNN1, MicroChemicals GmbH) attached to a standard 12.5mm stub with silver conductive ink (CAS: 7440-22-4, Alfa Aesar). Imaging parameters for ion beam focus and stigmator were determined by image optimization or autofunctions implemented in the microscope control software. To create the horizontal ion knife, a round feature such as a spot burn was used to increase the stigmator strength in horizontal direction through a combination of Stigmator X and Stigmator Y and refocus the feature into a line after the maximal stigmator strength was applied (see Supplementary Movie 1).

In arbitrary units within AutoScript4, the final stigmator values for imaging were - 0.027098 in x and 0.03531 in y. Working distance was at 0.01893 mm with the sample in coincidence point. For the ion knife, the stigmator values were 3 in x and -3 in y. Working distance was set to 0.01965 mm with the sample at coincidence point.

### 2.3 Beam shape characterization

Beam shaping experiments were performed on an Aquilos 2 and a Scios FIB-SEM system (Thermo Fisher Scientific). To measure the beam extensions at different currents, a piece of silicon (as in 2.2) was loaded into the FIB-SEM chamber and a spot burn was done for 10s. The imaging stigmator and defocus presets for ion knives and Gaussian-like probe were adjusted manually as described above. A 3×5 spot burn grid in x and y, respectively, was created automatically for all currents using a customized script in AutoScript4 (ThermoFisher scientific) version 4.3.1 with one row for horizontal, Gaussian-like, and vertical shapes (See Data and Code availability). Beam shapes were measured manually in FIJI [28].

For the volumetric beam shape analysis, a spot was milled at 30 kV and 1 nA and GIS coated using 8 kV 12 pA ion-beam assisted platinum deposition at a stage working distance of 6.8 mm. Images were collected in Auto Slice and View version 4.2.2 at pixel width of 2.7 nm (HFW 11.1 µm), 5 kV, 13 pA, 4096×3536 resolution and 1 µs dwell time for the shaped versus spot beam experiment and for the shaped versus scanned beam experiment 3.37 nm (HFW 5.2 µm), 5 kV, 25 pA, 1536×1024. Note that due to the changed GIS needle position on cryo-FIB instruments, filling was incomplete but still allowed for imaging and manually segmenting the probe. Images were aligned using SIFT in Fiji [29] and manually segmented using Napari. Segmentations were visualized using ChimeraX version 1.8 [30].

### 2.4 Lamella preparation

Lamella preparation was performed on an Aquilos 2 FIB-SEM operated at cryogenic temperatures < 180 °C. Spot burns were placed based on a previously collected image using the xT server user interface. Switching between probe and ion knife was done using custom scripts (see Code and Data availability). Exposures were done either manually moving the spot position towards the lamella or by placing line patterns (see Code and Data availability). The lamella was prepared using decreasing currents starting at 3 nA to 3 µm, 1 nA to 2 µm, 0.5 nA to 1 µm, 0.3 nA to 500 nm, and 0.1 nA to the final lamella thickness of ∼250 nm.

### 2.5 Lift-Out sectioning

Lamella preparation was performed on a high-pressure frozen *C. elegans* sample at cryogenic temperatures similar to protocols described before [24]. Platinum GIS deposition was done at a stage height of 6.8 mm for 60s. Trenches were milled using rectangular patterns at a current of 3 nA. Geometries of the lift-out volume were circa 40 µm x 40 µm in x and y, respectively. Sample thickness is determined by the ‘waffle’ protocol used and is around 25 µm. Attachment using a copper block adaptor was performed as described before [24]. After transfer and alignment to the 100/400 mesh copper grid bars, the sectioning step was performed using a stigmatic beam at 3 nA for a sectioning thickness of 1 µm and 1 nA for a sectioning thickness of ∼300-400 nm. To align the probe with the block to be sectioned, exposures were first done on the grid bars and the spot was subsequently manually moved towards the lift-out volume’s front.

### 2.6 TEM data acquisition

TEM data was acquired on a Titan Krios G2 equipped with a XFEG, K2 camera and a GIF energy filter at 42 000 x nominal magnification resulting in a pixel size of 0.342 nm at the sample. Tilt series were acquired using SerialEM version 4.1.0 beta [31]. Tilt range was from -70° to +50° starting at the lamella pre-tilt of 10° and using a dose-symmetric tilt scheme [32]. Total fluence corresponds to 120 e A-2. The frames were aligned in motioncorr2 version 1.4.7 [33] and reconstructed at bin4 using 3DMOD version 4.11.15 [34] and SIRT-like filter at 20 iterations.

## 3. Results

### 3.1 Generation of ion knives using the FIB stigmator

Pursuing the idea of shaping the ion beam, we explored to what extent the readily available stigmator could be used to generate a line-shaped beam from the ion optics present in modern FIB-SEM instruments (Figure 1a). In standard operation, the slightly astigmatic beam is corrected by the stigmator to generate a rotationally symmetric or Gaussian-like beam profile (Figure 1b, standard usecase). In the proposed approach, we aim to increase the stigmator strength in the horizontal direction to the maximum, generating an ion knife (Figure 1b, knife usecase). To that end, we used an in-house python-simulation to explore the generation of highly astigmatic beams from a Gaussian-like beam profile (Figure 1c). In the vectorized model of an electrostatic quadrupole, the Gaussian-like beam was virtually imaged after a quadrupole stigmator with varying strengths, simulating the conventional use case of a circular probe with a low stigmator strength and the new use case of ion knife with high stigmator strength, where the beam is distorted towards a slim, horizontal ellipse.

**Figure 1:**
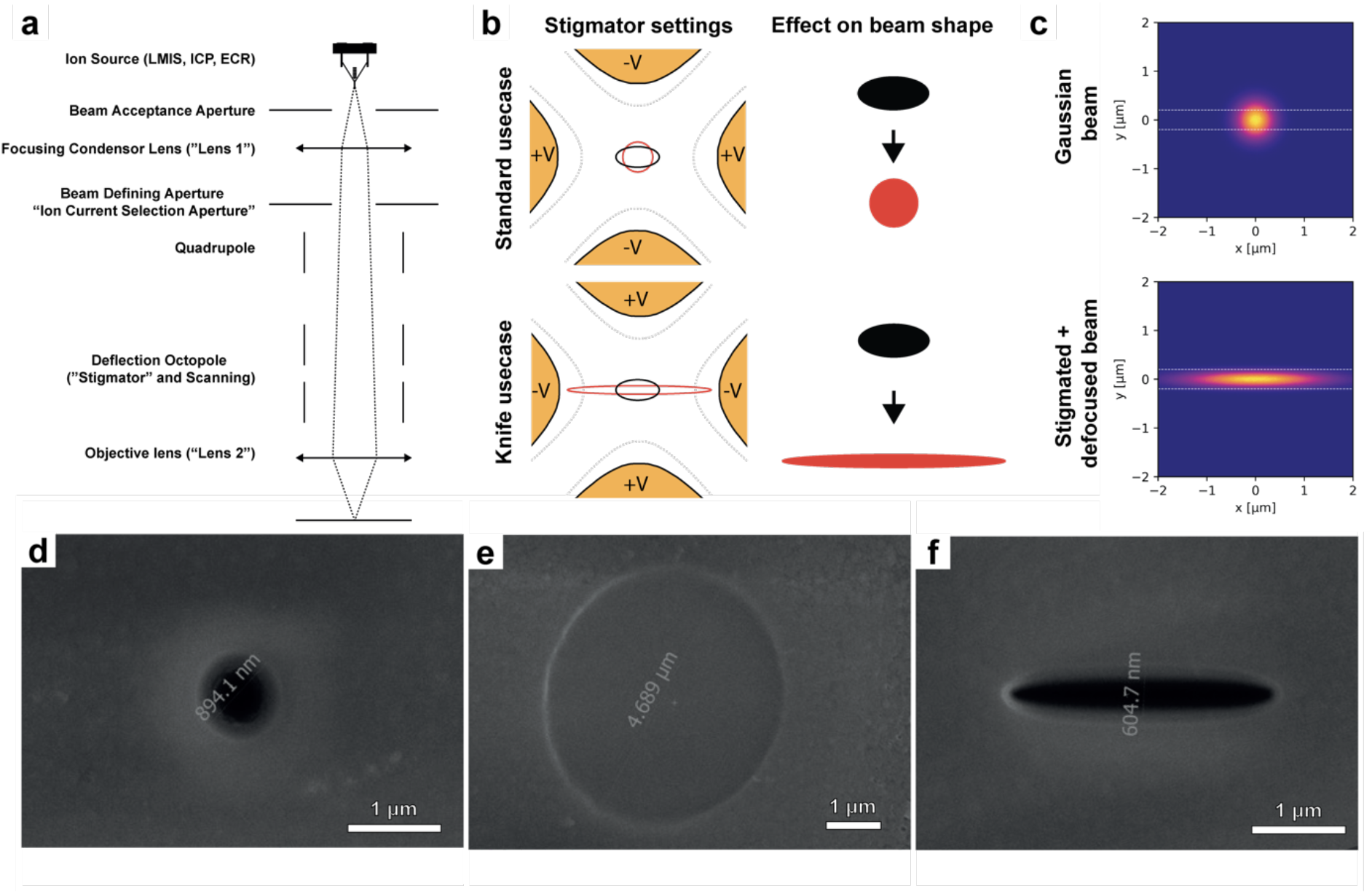
Generating ion knife shapes using the FIB stigmator and objective lens. **a**, Schematic of an ion column. **b**, Schematic representation of the stigmator and its qualitative effect on the beam shape. Black shapes represent the beam profile prior to stigmation, red the beam profiles with adjusted stigmator values. **c**, Simulation of the effect of stigmation on the shape of a Gaussian-like beam profile. Starting with a Gaussian-like ion beam profile (top), astigmatism in X direction is applied leading to a stretching of the Gaussian-like beam in one direction (bottom). White dashed lines at a constant position for reference. **d**, Spot burn for 10s at 1 nA using a stigmatic, focused probe. **e**, Spot burn for 10s at 1 nA using an ion beam probe with applied stigmator strength. **f**, Spot burn for 10s at 1 nA using an ion beam probe with applied stigmator strength and adjusted focus.

Stigmation alone is not sufficient to spread the ion beam to create a line shape as it will mostly generate a blurred beam in reality. Increasing the stigmator strength in a specific direction can be understood as creating a lens with two different focal points. Therefore, when defocusing a highly astigmatic beam, the focal point in one direction, e.g. the vertical direction, is approached while moving away from the focal point in the orthogonal (horizontal) direction. This leads to an elongation of the probe profile in one direction and sharpening of the probe profiling in the perpendicular direction.

In practice, shaping the ion beam to a knife can be performed by the following steps: First, the ion beam in normal ‘imaging’ mode is focused and stigmated as in standard operation. This leads to a spot burn that resembles a Gaussian-like profile (Figure 1d). Subsequently, the stigmator values are driven to the extreme. As the stigmator multipoles are neither mounted parallel nor perpendicular to the horizontal axis, a combination of both the X and Y stigmator values is needed to create the horizontally-oriented shape of the ion beam. As this stigmation alone will only produce a highly defocused beam, as described above, a spot burn at this stage results in a large circle (Figure 1e), resembling the circle of least confusion of the astigmatic beam. Defocusing the beam will bring it closer to the focal point in the vertical direction. Therefore, the beam diameter of the spot burn decreases in the vertical direction while the beam width increases in the horizontal direction (Figure 1f). The resulting spot burn shows an anisotropic resolution. Seemingly, the ion beam shape becomes narrower in height, creating the beam that we term ‘ion knife’. A practical example of creating the ion knife is best illustrated on a single bright or dark spot on the sample, e.g. a spot burn (Supplementary Movie 1).

### 3.2 Shape characterization of ion knives

The creation of spot burns and their subsequent imaging or even cross-sectioning is a relatively time-consuming method to determine the shape of a probe. In order to identify optimal parameters of beam shapes in a rapid manner, we developed a method to approximate the beam shape through imaging rather than through spot burns and subsequent cross-sections in analogy to the previous described generation of the knife (Supplementary Movie 1). In order to measure the approximate beam shape via imaging (Figure 2a), we create a spot burn with a Gaussian-like probe using the standard beam settings (Figure 2b). Subsequently, the spot burn is imaged with the Gaussian-like standard (Figure 2b, left) and the shaped probe (Figure 2b, right), resulting in an image that represents the convolution of the spot burn with the shaped ion beam (Figure 2a). Comparing the probe measurement via imaging (Figure 2b) to the measurement via spot burn (Figure 2c) shows good agreement between beam profiles in Gaussian-like probe as well as in ion knife mode. Moving the astigmatic probe through focus leads to sharpened beam profiles in the horizontal (−Defocus) or vertical (+Defocus) direction as described above (Figure 2d), effectively rotating the ion knife when going through the focal point. In order to screen stigmator parameters to find optimal beam shapes for the application of a horizontally aligned ion knife, we generated two-dimensional parameter matrices as a beam shape tableau (Figure 2e).

**Figure 2:**
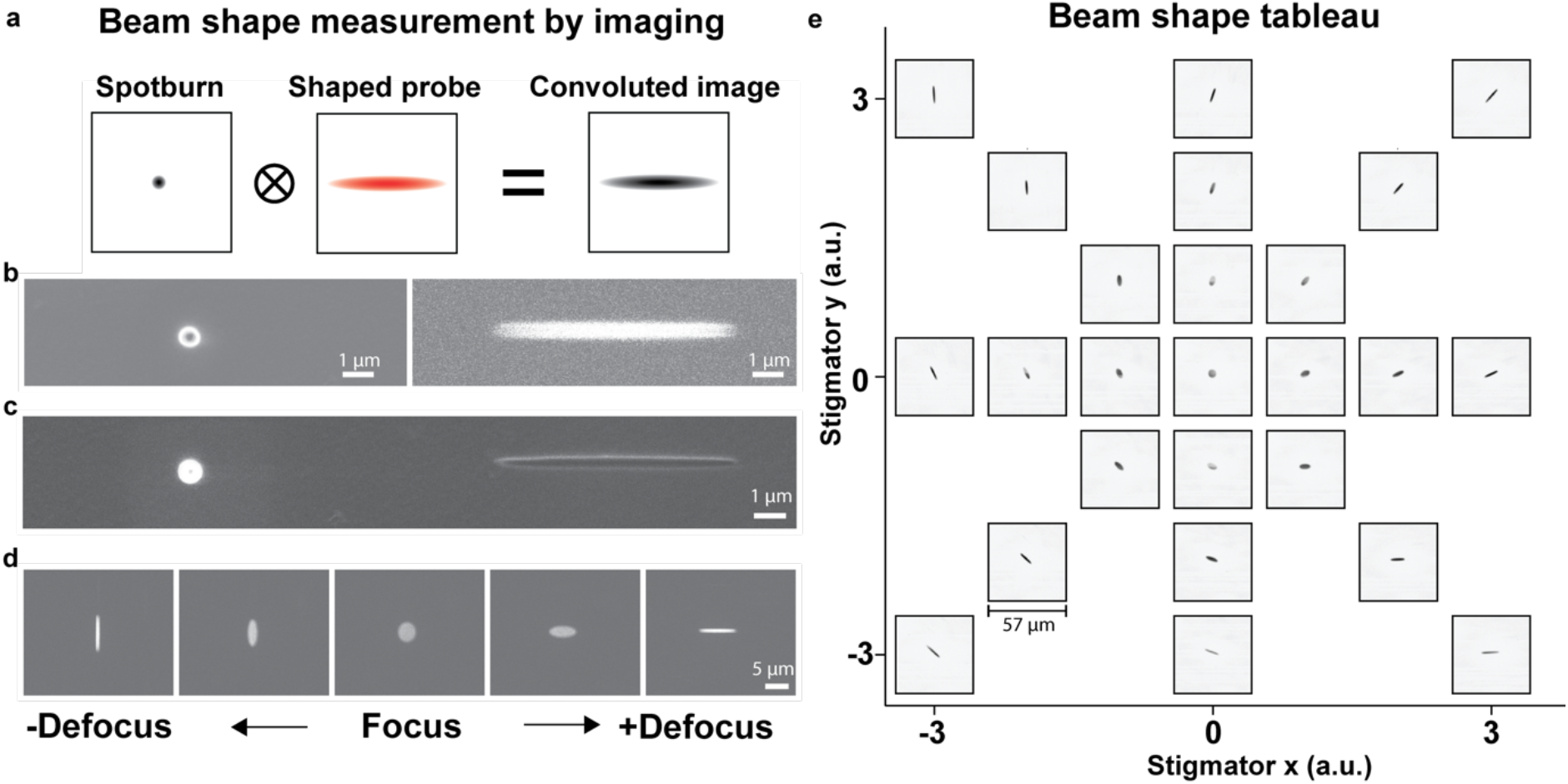
Beam shape measurement by imaging. **a**, Principle of the imaging-based approach for parameter screening of the ion knife. A spot burn is imaged with a shaped probe, resulting in a convoluted image that resembles the shape of the probe. **b**, Spot burn with a Gaussian-like beam profile (left) imaged with a knife shaped probe (right). **c**, Side-by-side comparison of a spot burn using a Gaussian-like probe (left) with a spot burn using a knife-shaped probe (right). **d**, Defocusing the probe will yield a sharpened probe that changes orientation by 90 degrees comparing the under- and overfocused profile. **e**, Assembling the beam shape images of the stigmator values into a matrix or beam shape tableau allows to find the ideal parameters for shaping the probe.

To quantify whether the shaping of the beam diameter and length is achievable across beam defining apertures, we characterized spot burns of ion knives and Gaussian-like probes at different positions of the beam-defining aperture strip, i.e. at different currents. The shape of an ion knife burn is defined by two different parameters, its length and its width or minimal diameter (Figure 3a), while an ideal spot burn using a Gaussian-like probe is defined by a single parameter, its diameter (Figure 3a). If the stigmator strength is increased to its maximum, sharp lines can be achieved during ion beam milling (Figure 3b). By exposing multiple positions on a silicon sample for a defined time of 10s while switching between horizontal knife burn, Gaussian-like probe burn, and vertical knife burn, we determined these shape parameters. As only electrostatic elements are switched back and forth from knife and spot optics during this process, rapid switching between imaging and knife mode is feasible and hysteresis is negligible in the process. These characterizations optimized stigmator parameters for horizontal ion knives, as they show the highest potential for TEM lamella milling. The same parameters with reversed sign were used to create the vertical ion knife beam profile, thus leading to slightly sharper horizontal compared to vertical shapes (Figure 3b).

**Figure 3:**
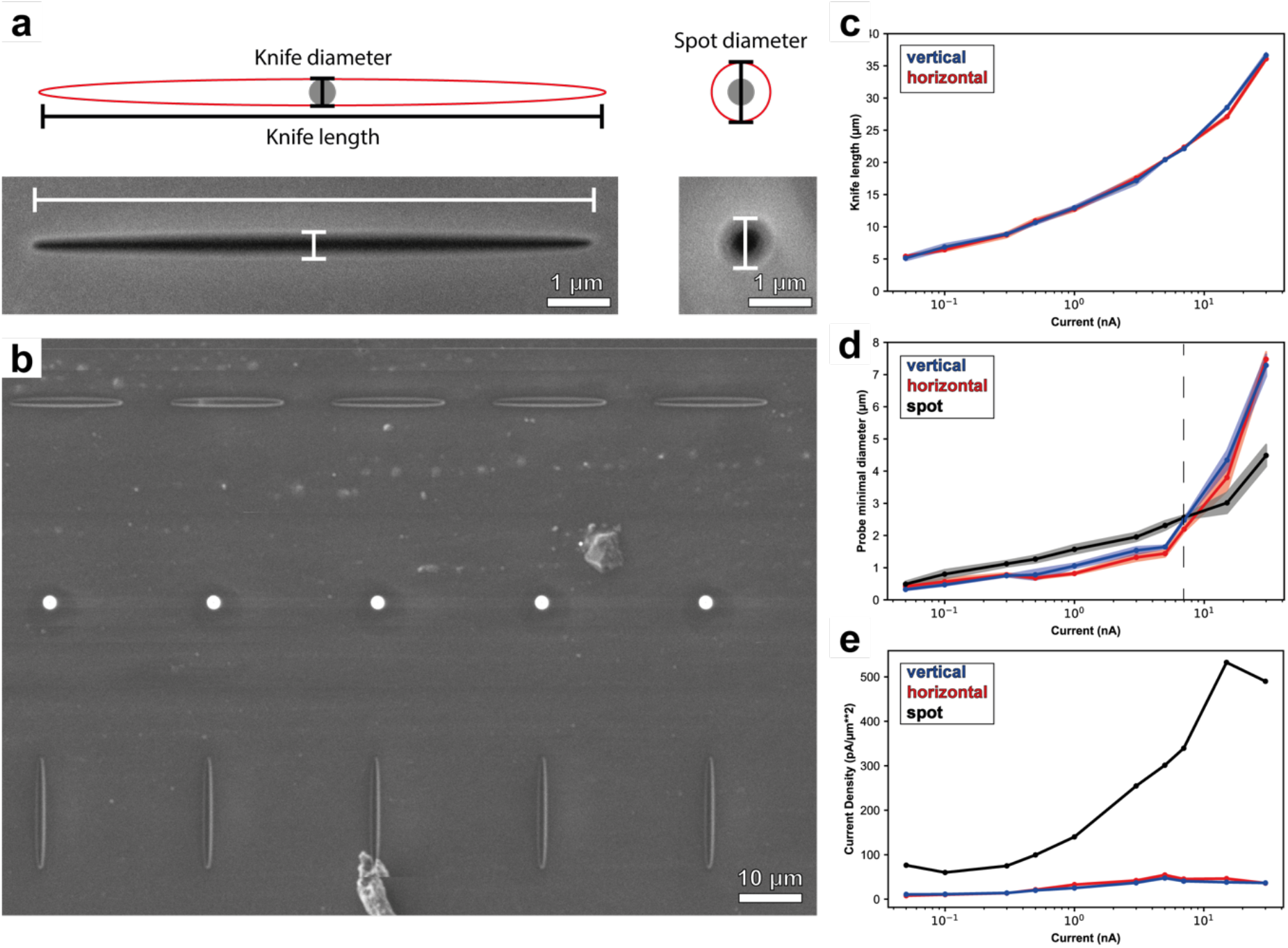
Shape characterization of ion knives using spot burns. **a**, Schematic of the measured beam parameters of the ion knife and the Gaussian-like ion probe. The ion knife geometry is described by its length and diameter, while the Gaussian-like ion probe is described only by its diameter. **b**, Example image of spot burns using horizontal knife, Gaussian-like, and vertical knife (top to bottom) ion beam profiles. Spot burns were performed switching back and forth between ion probe shapes. c-e, probe characterization for vertical ion knife (blue), horizontal (red) ion knife and Gaussian-like spot (black) probes **c**, Ion knife lengths across different currents. **d**, Minimal diameter of spot burns using Gaussian-like probes and ion knives in vertical and horizontal set up. **e**, Current density of Gaussian-like ion probes and ion knives assuming uniform dose distribution across the beam profile.

The knife length is determined by the current given by the beam defining aperture size as the stigmator strength is constantly at its maximum. It ranged from 5,1 µm ± 0,4 µm at 50 pA to 36,6 µm ± 0,5 µm at 30 nA (Figure 3c, Supplementary Figure 1). As expected from the simulations (Figure 1c), the ion knife diameter judged by silicon burns changed compared to the Gaussian-like probe diameter (Figure 3d). For currents commonly used in TEM lamella preparation (50 pA – 7 nA), the ion knife shows consistently decreased minimal diameter than the Gaussian-like probe diameter at the same current (Figure 3d). As the stigmator and focus parameters were optimized for horizontal ion knife shaping, the minimal knife edge in the horizontal direction was also consistently smaller than in the vertical direction. Assuming uniform distribution of the current across the FIB probe, the current density increases when using a Gaussian-like probe with ∼ 70 pA/µm^2^ at 50 pA and ∼350 pA/µm^2^ at 7 nA while showing a lower current density when using the ion knife with ∼10 pA/µm^2^ at 50 pA and ∼50 pA/µm^2^ at 7 nA (Figure 3e). The current density of the ion knife seems to be practically limited to ∼50 pA/µm^2^.

### 3.3 Beam profile comparison

To evaluate the beam depth profile of spot burns and ion knives, we created cross sections of spot burns of different positions of an ion knife (Figure 4a) and of a Gaussian-like probe (Figure 4b). Despite our initial intuition to expect drastic variations in depth and shape for cross-section profiles along the ion knife, the experimental results show a relatively uniform depth distribution. The profile of the ion knife only differs marginally going from the center to the extreme of the knife (Figure 4c-e, g), with a milling depth difference of ∼200 nm along the length axis for a 10s exposure. However, the comparison to the spot burn of a Gaussian-like probe (Figure 4f) shows that the drop in current density of the ion knife leads to a decrease in factual milling depth to only around 30% of the spot burn. Interestingly, at similar distance to the deepest point of the milling trench, the shape of the ion knife shows a narrower profile compared to the Gaussian-like probe (Figure 4g). Using FIB-SEM tomography on exposures of Gaussian-like probe and knife probe beam shapes on silicon, we determined the 3D shape of a spot burn versus a knife burn (Figure 4h-i, Supplementary Movie 2, Supplementary Movie 3). This allowed us to approximate the value of the volume of ablated material in silicon of a 10 s spot versus knife burn at 1 nA, namely 0,38 µm^3^/s for a spot burn and 3,32 µm^3^/s for the ion knife. Comparing cross-sections of a shaped ion knife burn (Figure 4j,l) to a line-scanned Gaussian-like probe (Figure 4k,m), we observe an approximately 2x sharper burn for the ion knife (Figure 4n) while the rate of volume ablation remains the similar, with 0,42 µm^3^/s and 0,39 µm^3^/s for scanned probe and ion knife at 0.3 nA, respectively.

**Figure 4:**
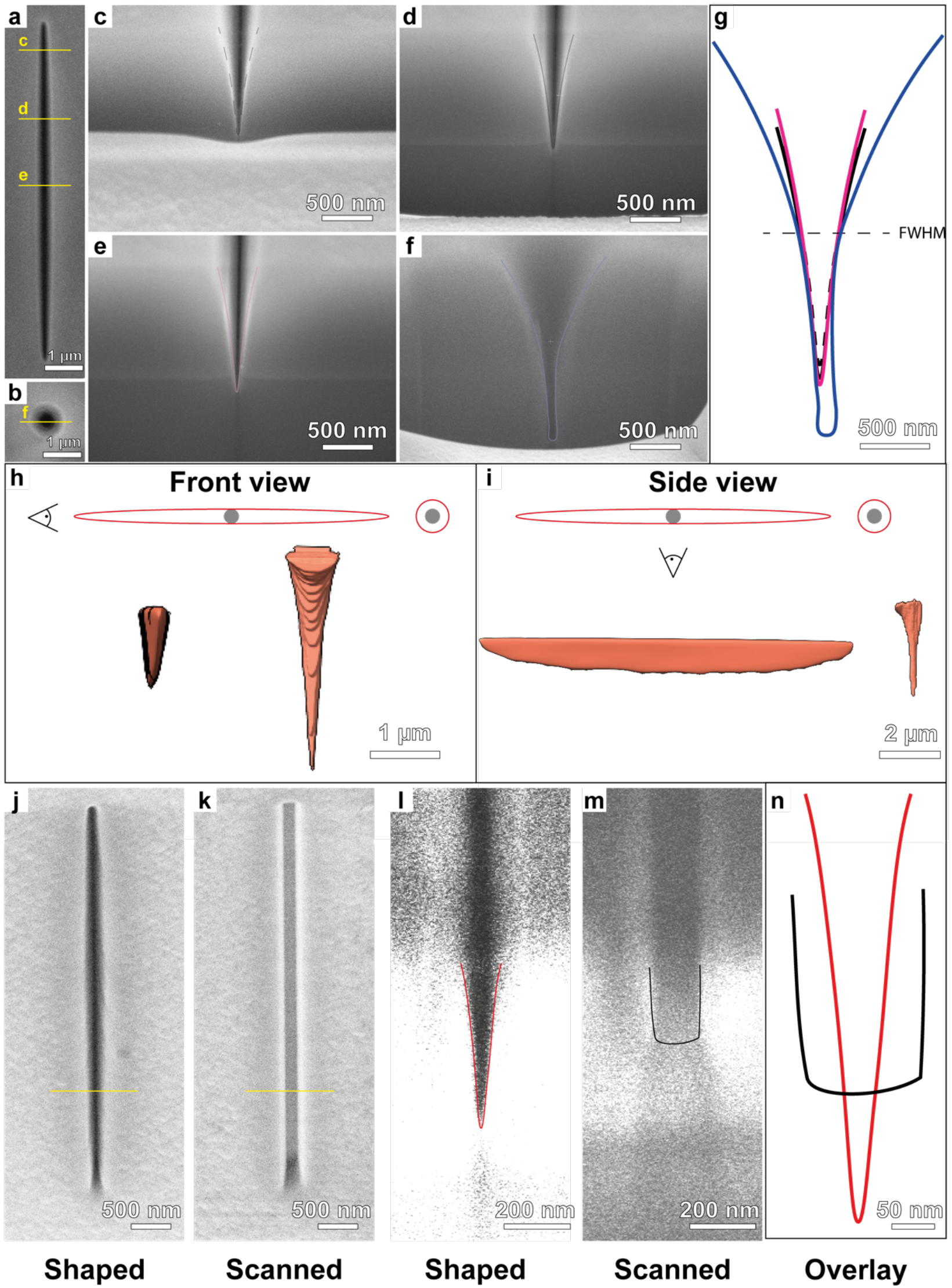
Comparison of beam shapes along the ion knife versus the standard probe shape and scanned versus shape probe spot burns. **a**, Ion beam spot burn using an ion knife. Cross-section profile positions of the ion knife at different positions are indicated. Profiles are shown in panels c-e. **b**, Ion beam spot burn using a Gaussian-like probe. Cross section of the spot burn shown in f. **c-e**, Cross-section of the ion knife at positions indicated in a. **f**, Cross-section profile of the Gaussian-like ion beam spot burn shown in b. **g**, Overlay of the measured ion beam profiles. Gaussian-like spot burn shown in blue, profiles from c in dashed black line, d in solid black line, and e in solid magenta line. Beam profiles are aligned to the full width at half maximum (FWHM). **h**, Front and **i**, Side view of the 3D rendering of spot and knife burns obtained by FIB SEM tomography. **j**, FIB image of an ion knife spot burn after 10s exposure. Yellow line indicates the cross-section position in l. **k**, FIB image of a line-scanned Gaussian-like probe after 10s exposure. Yellow line indicates the cross-section position in m. **l**, SEM image of a cross-section of the ion knife burn in j. **m**, SEM image of a cross-section of the line-scanned Gaussian-like probe burn in k. **n**, Overlay of the beam profile cross-sections from l and m. Black profile corresponds to the scanned Gaussian-like probe, red represents the ion knife.

### 3.4 Milling cellular lamellae with ion knives

To evaluate the feasibility of using ion knives for cryo-ET sample preparation, we deployed the information obtained on probe shaping to apply ion knife milling to the generation of lamellae on a TEM grid from cells vitrified by plunge freezing (Figure 5a). By exposing positions with the ion knife along a vertical line approaching the final lamella position in a sequential fashion of decreasing current (Figure 5b), we were able to prepare lamellae from clumps of cells on the grid (Figure 5c-d). The narrowing of the lamella towards its final width, which is commonly done in on-grid cryo-lamella workflows for cellular samples, is a direct result of the narrowing beam shape as one moves to smaller currents or beam-defining aperture positions. However, due to the limited length of the beam at low currents, the imageable region of the lamella becomes relatively small, < 8 µm. Multiple positions along the width of the lamella can be exposed to create a larger lamella, however in our experiments this led to a drastic increase in curtaining (Supplementary figure 2). The preparation time for lamellae prepared with the ion knife was ∼15-20 minutes, comparable to normal lamella preparation. With improved software integration, the throughput could definitely be improved as the current protocol is tedious and mostly manual work (Figure 2e). Notably, due to the decreased current density, polishing was performed at 300 pA, which also shows curtaining during milling with standard Gaussian-like FIB probes. Despite the limited size of the lamella when using single beam exposure positions, we were able to collect tilt series in the cryo-TEM on these lamellae (Figure 5e). The reconstructed tomogram of a starved yeast cell shows the expected features such as large stress granules and ribosomes within the crowded cellular environment (Figure 5f).

**Figure 5:**
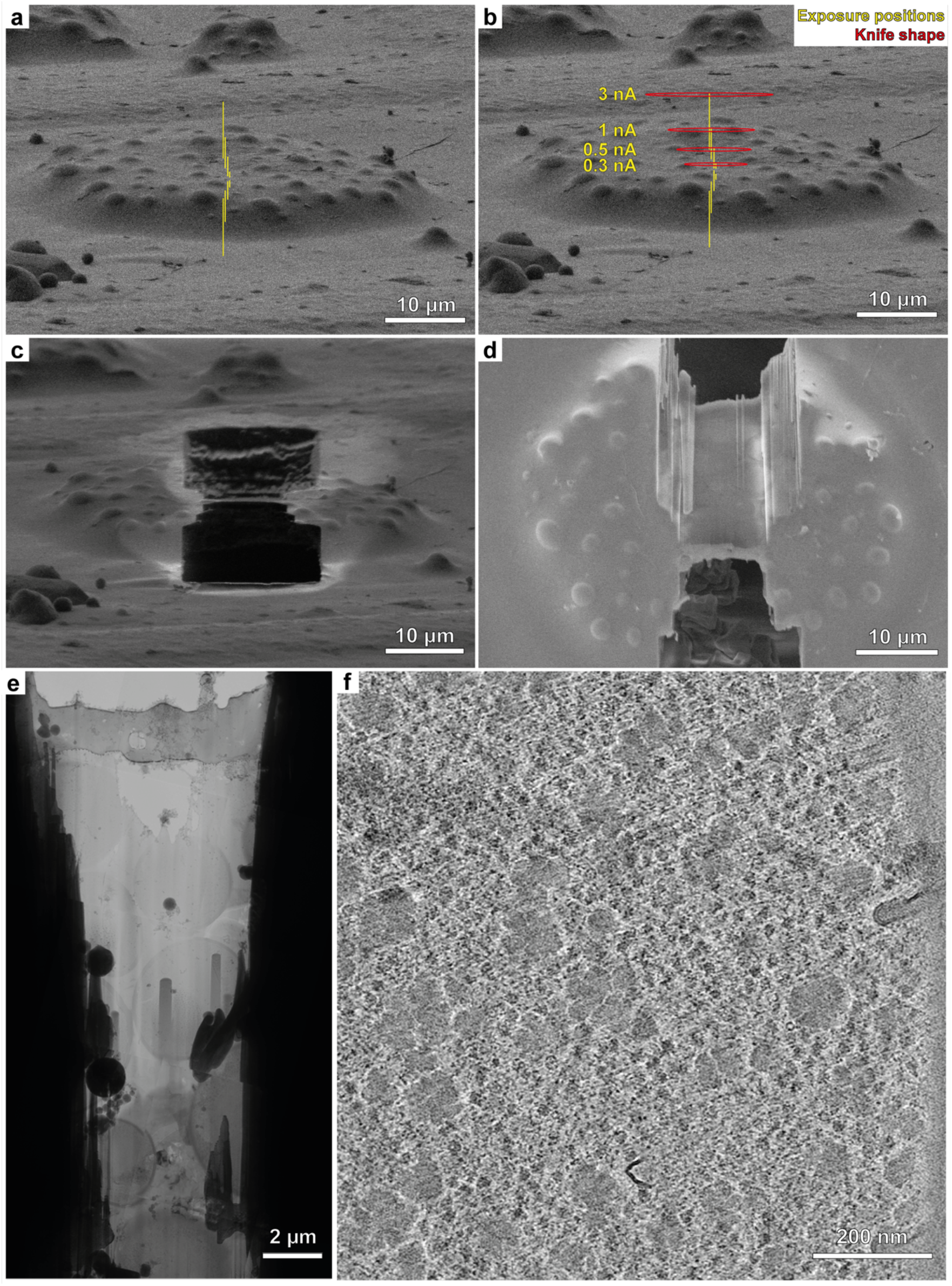
Preparation and cryo-ET imaging of ion knife milled lamellae. **a**, FIB image of the cells prior to exposure. **b**, Schematic of the lamella milling process using ion knives. Yellow lines in a and b indicate the exposure positions, red ellipsoids the approximate shape of the beam used for milling. **c**, FIB and **d**, SEM view of the prepared lamella. **e**, TEM overview of a lamella milled with the ion knife. Note that the viewing window is limited as only single positions were exposed, leading to decreasing lamella width with decreasing current. **f**, Slice through a reconstructed tomogram from a tilt series collected on a lamella milled with ion knives. Slice thickness is 1.4 nm.

### 3.5 Using ion knives for sectioning in Serial Lift-Out

Lamella preparation workflows for on-grid lamellae are established and have become a widespread application in specialized cryo-ET laboratories across the globe. Therefore, the gain from ion knives could be a potential increase in throughput if specific hardware and software integration is developed. Another application where we see the benefits of ion knives becoming useful is in the generation of vitreous lamellae from cryo-lift-out specimens using the Serial Lift-Out method. Here, the ion knife holds the potential to reduce the material loss in sectioning due to the sharpening of the probe in the vertical direction as well as potentially circumvent often observed redeposition of the sectioned lamella that prevents its detachment from the bulk lift-out material. To that end, we performed a proof-of-concept experiment using the Serial Lift-Out method (Figure 6, Supplementary figure 3). In Serial Lift-Out, a large volume is extracted from the specimen using a micromanipulator needle (Figure 6a-d) and then sliced into thinner sections, usually around 1-3 µm, which are thinned to less than 200 nm for cryo-ET [24]. In conventional milling approaches, more than 500 nm of the lift-out material is lost between the sections. Combined with the thinning process of the thick sliced sections, this leads to a large loss of material in the ‘quasi-sequential’ lift-out sectioning. Therefore, it is desirable to minimize the material loss as much as possible. In order to demonstrate sectioning with the ion knife, we used an ion knife at 1 nA for lamella sectioning during the Serial Lift-out process (Figure 6e-h). While the ion knife remains complicated in its usage, slicing a section from the lift-out block at an apparent thickness of 300-400 nm as judged from the FIB image was achievable (Figure 6g-h).

**Figure 6:**
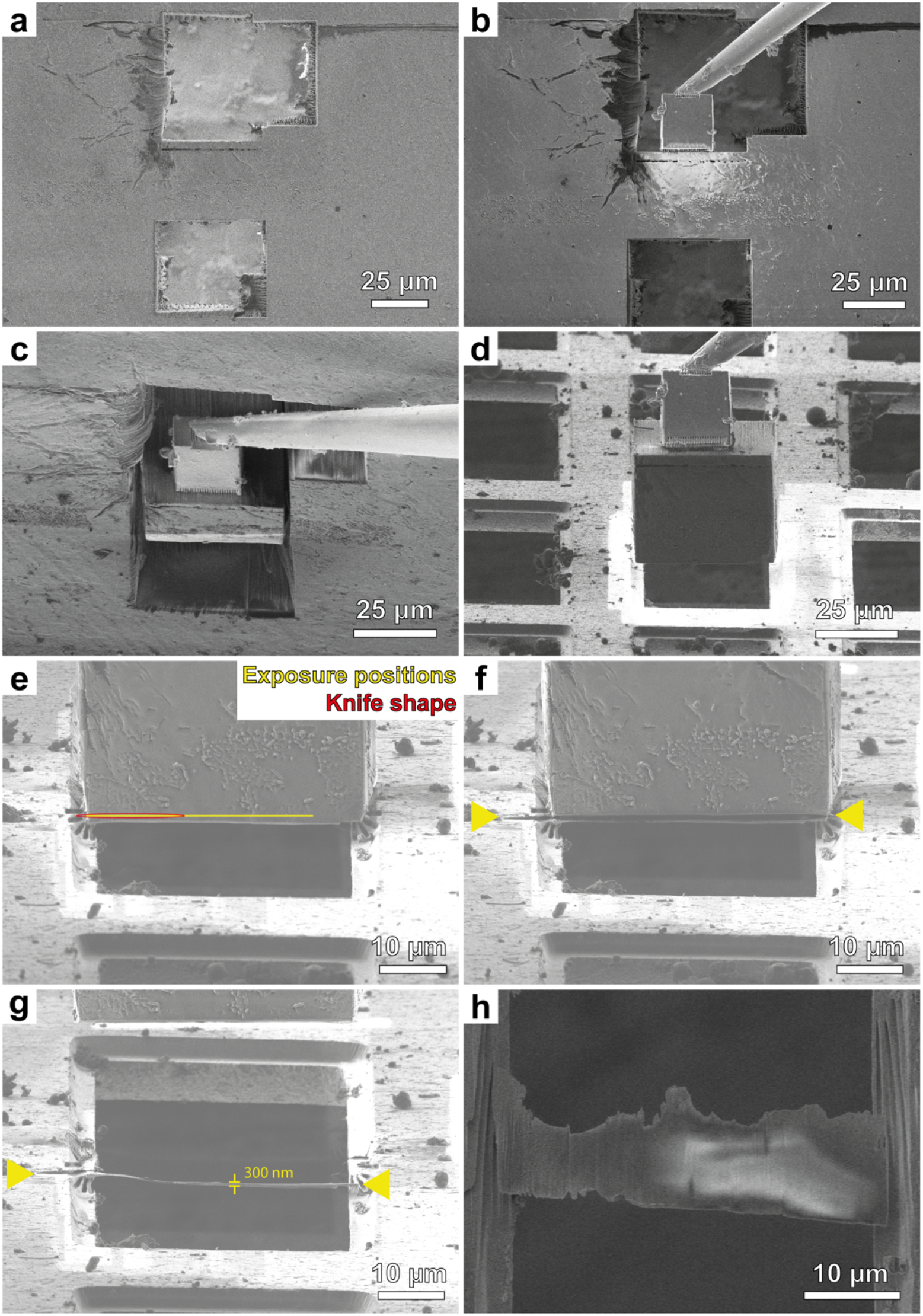
Pilot experiment of Serial Lift-Out sectioning with the ion knife. **a**, FIB image of vitrified material after trench milling. **b**, The volume of interest is approached with the copper block adaptor on the micromanipulator and attached via redeposition milling at the interface **c**, The volume of interest is dispatched from the bulk by FIB milling and extracted using the micromanipulator. **d**, The receiver grid, here a 100/400 mesh copper grid, is approached. **e**, The block is aligned to the receiver grid and the lower region of the lift-out volume is attached to the receiver grid by redeposition milling patterns. The red oval shows the beam profile, the yellow line corresponds to the beam exposure positions. **f**, FIB image after exposure with the ion knife. Yellow arrowheads indicate the milled regions. **g**, FIB and **h**, SEM image of the lamella formed by the ion knife after the remaining extracted volume is lifted up.

## 4. Discussion

In pursuing the idea of shaped FIB probes in lamella preparation, we demonstrate the generation and usage of highly anisotropic but defined beam shapes in the FIB instrument by using the available stigmator and the objective lens (Figure 1a, “lens 2”) of the ion column. We characterize the shape of the generated beam profiles that we term ‘ion knives’ by silicon spot burns and FIB-SEM volume imaging. We explore its potential applications in cryogenic lamella preparation of cellular material and the Serial Lift-Out protocol for high-pressure frozen samples. Through our characterization, we find that the dimension of these ion knives is given by the beam defining aperture. Furthermore, the switching between imaging using Gaussian-like probes and milling using ion knives is feasible when using the stigmator as a quasi-cylindrical lens. While we have only characterized horizontal and vertical orientation of the ion knife, arbitrary angles can be selected by choosing the X and Y stigmator strength accordingly. It needs to be noted that a widely adopted alternative method to measure beam performance in the FIB is the knife edge method. However, as the method requires the measurement of an image on a sharp edge, it is difficult to measure beam performance with the highly astigmatic beam of the ion knife as no real image can be produced. As an alternative to the measurement of spot burns, we introduce the concept of probe shape measurement by imaging. We introduce the beam shape tableau that allows the rapid determination of approximate beam shapes via the imaging of a Gaussian-like spot burn. This technique is especially useful when adjusting parameters such as stigmator values or aligning mechanical parts where quick feedback is critical. While it was primarily used here for configuring the stigmator and defocus, the method has since helped us optimize other parameters (e.g. condenser strength C1).

To characterize beam shapes at higher resolution, we determine example beam shapes through measurement of cross sections of spot burns in silicon. In contrast to our initial hypothesis, the milling depth along an ion knife burn remained relatively constant (Figure 2a,c-e), which is beneficial for applications in lamella preparation. Furthermore, our experiments suggest that for the same current, ion knives ablate a greater volume of material from a surface at the cost of milling depth than standard FIB spots (Figure 2a-i). However, we did not observe these differences when comparing a stationary ion knife with a scanned Gaussian-like probe exposure. We thus conclude that the observed effect is a result of the limited milling depth for ion knife burns. The reduced milling depth is likely the result of a limited current density using ion knife shapes. Currently, this is detrimental to the lamella milling workflow as the limited current density and primitive scanning strategy tends to give increased curtaining effects. This could, however, be overcome with specifically designed hardware, e.g. shaped apertures, a true cylindrical lens in the ion beam path instead of using the stigmator multipoles, and a better software integration. Nevertheless, as the whole milling area besides the knife ends is continuously exposed to the ion beam, we speculate that with adjusted designs redeposition of ablated material could be diminished.

As commonly available apertures in modern FIB instruments are circular with increasing diameter for higher currents, the beam’s width when spread to an ion knife is dictated by the aperture size: the higher the current, the wider the knife dimension will be. The current density seems to almost remain constant when changing aperture size (Figure 3e). This could potentially be mitigated by adjustments in hardware, for example by shaping the aperture, e.g. to a slit, or as mentioned by developing dedicated cylindrical lenses. Since plasma FIB systems are becoming increasingly available to the cryo-ET community [10, 11, 35, 36], it is worth noting that when moving to inductively coupled plasma FIBs, initial experiments suggest that the beam shapes become even larger than when using liquid metal ion sources for the currents used for biological lamella preparation. This is likely due to a combination of a much larger virtual source size for the ICP (the PFIB source available to us) compared to LMIS that leads to beam tails in the spot burn profile [9] and differences in aperture sizes of the beam determining apertures going from LMIS FIB to ICP PFIB columns.

Biological processes happen in context, meaning that the surrounding material of a lamella is of utter importance. Consequently, it is desirable to maximize the portion of the initial sample that becomes imageable volume in the TEM. With standard on-grid lamella milling and lift-out approaches at cryogenic temperatures, only a fraction of the specimen (usually <1% for eukaryotic cells and orders of magnitude less for multicellular organisms and tissues) ends up in the TEM ready for cryo-ET data acquisition. One way to mitigate this, recently described by multiple labs [23, 24], is the approach of milling multiple lamellae from a single lift-out volume. In Serial Lift-Out, the material loss is determined by the sectioning thickness and the amount of lost material due to the ablation process as a lamella is sliced from the large lift-out volume. By sharpening the ion beam profile as demonstrated here, the lost material can potentially be reduced. Additionally, our pilot experiments suggest that thin sections <400-500 nm can be obtained directly from the block using the ion knife. While we consider it an interesting proof-of-principle, improvements in both hardware and software will be necessary to make this method adaption available to the FIB practitioner. Especially the software integration will be a challenge of its own, requiring calibrations and alignment routines for the shifts between imaging spots and knife spots, as well as adjusted focusing routines for the ion knife in order to get the best possible probe. However, if these technical and technological shortcomings are addressed, the method holds the potential to enable milling with higher throughput as more current can be spread over larger areas as well as smaller material loss during Serial Lift-Out. We anticipate that FIB shaping will allow for more contiguous volumes from biological specimens, bringing lift-out sectioning a bit closer to the established serial sectioning with diamond knife approaches in cryo-electron microscopy of vitreous sections (CEMOVIS) [37, 38] while maintaining the level of structural preservation that cryo-FIB milling offers. In conjunction with developments in the acquisition of large areas on lamellae [39-43], this would enable tracing large fractions of entire organelles and cells in cryo-ET, generating an increasingly holistic view on cellular ultrastructure. While substantial development will be required, FIB shaping could bring us a little closer to the structural cell biologist’s dream: quasi-continuous reconstructions of entire multicellular organisms at molecular resolution.

## Data and code availability

All data will be made available upon request. To enable others to experiment with the ion knife, we developed jupyter-notebooks in which pre-stored values for horizontal and vertical ion knives are loaded and manually refined. Code used throughout the work to form ion knives as well as pattern files to mill cells are available on GitHub: https://github.com/sklumpe/beamshape-manuscript.

## Author contributions

JB and SK performed experiments, SK and JB conceptualized experiments, JMP and SK supervised the work, SK wrote the initial draft of the manuscript with input from all authors. JB, JMP, and SK edited the manuscript. JMP and SK acquired funding.

## Acknowledgements

We would like to thank Thomas Juffmann, Joakim Reuteler, Stephan Gerstel, and David Klebl for fruitful discussions on beam shaping. We thank Christoph J.O. Kaiser & Oda H. Schiøtz for providing HPF samples. We thank the members of the Plitzko and Briggs labs for insightful discussions. This study used the infrastructure of the Department of Cell and Virus Structure at the MPI of Biochemistry. JMP acknowledges funding from the Max-Planck Society. SK acknowledges the support and the use of resources of Instruct-ERIC through the R&D pilot scheme APPID 3485.

