## Supplementary Information for "Exploring shaped focused ion beams for lamella preparation"

**Supplementary movie 1: Demonstration of ion knife generation using the beam shape measurement by imaging method.** A previous spot burn is imaged at 30 kV and 1 nA, which gives a first approximation of the beam shape used to scan the sample for imaging. While imaging, the stigmator is increased in one direction, leading first to a defocused beam. After focusing, this beam becomes a sharper, astigmatic beam. Finally, settings are changed back to default stigmator values and the spot is refocused.

**Supplementary movie 2: FIB-SEM tomography of a 10s exposure in silicon using a Gaussian-like profile beam.**

**Supplementary movie 3: FIB-SEM tomography of a 10s exposure in silicon using a shaped ion knife beam.**

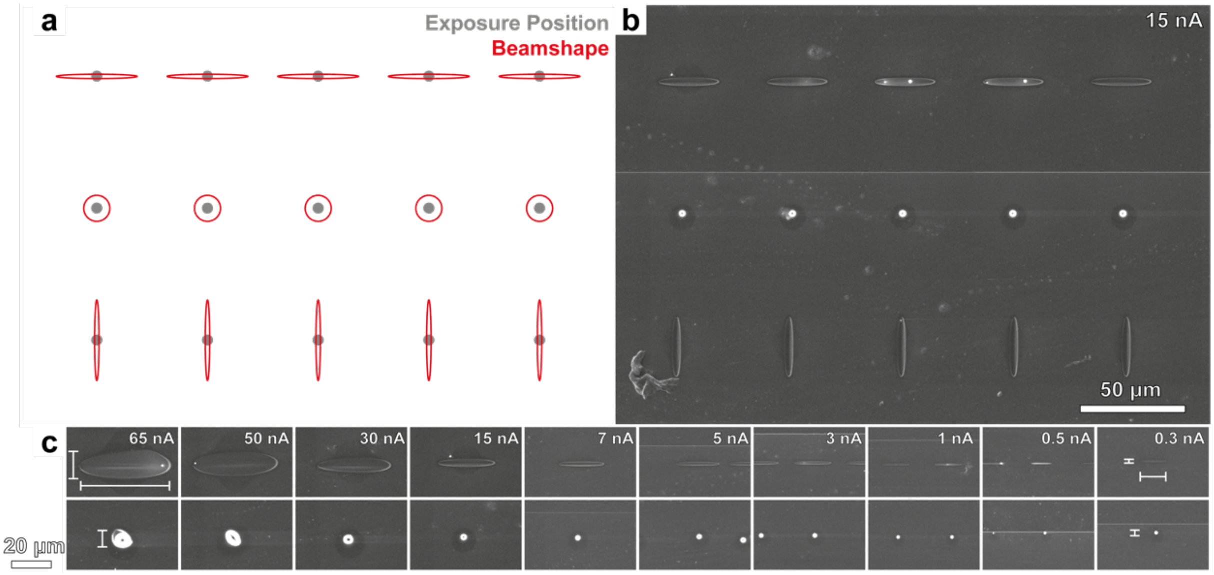


**Supplementary figure 1: Beam Shape examples for different currents. a,** Schematic of the beam shape experiment. Shown are the exposure points and beam shapes. A 5x3 grid is creating with the first row representing horizontal ion knives, the second-row Gaussian-like beam profile spot burns, and the third-row vertical ion knives. **b,** Spot burn example for the 15 nA beam determining aperture position. **c,** Horizontal ion knives and spherical beam profiles in side-by-side comparison across beam currents.

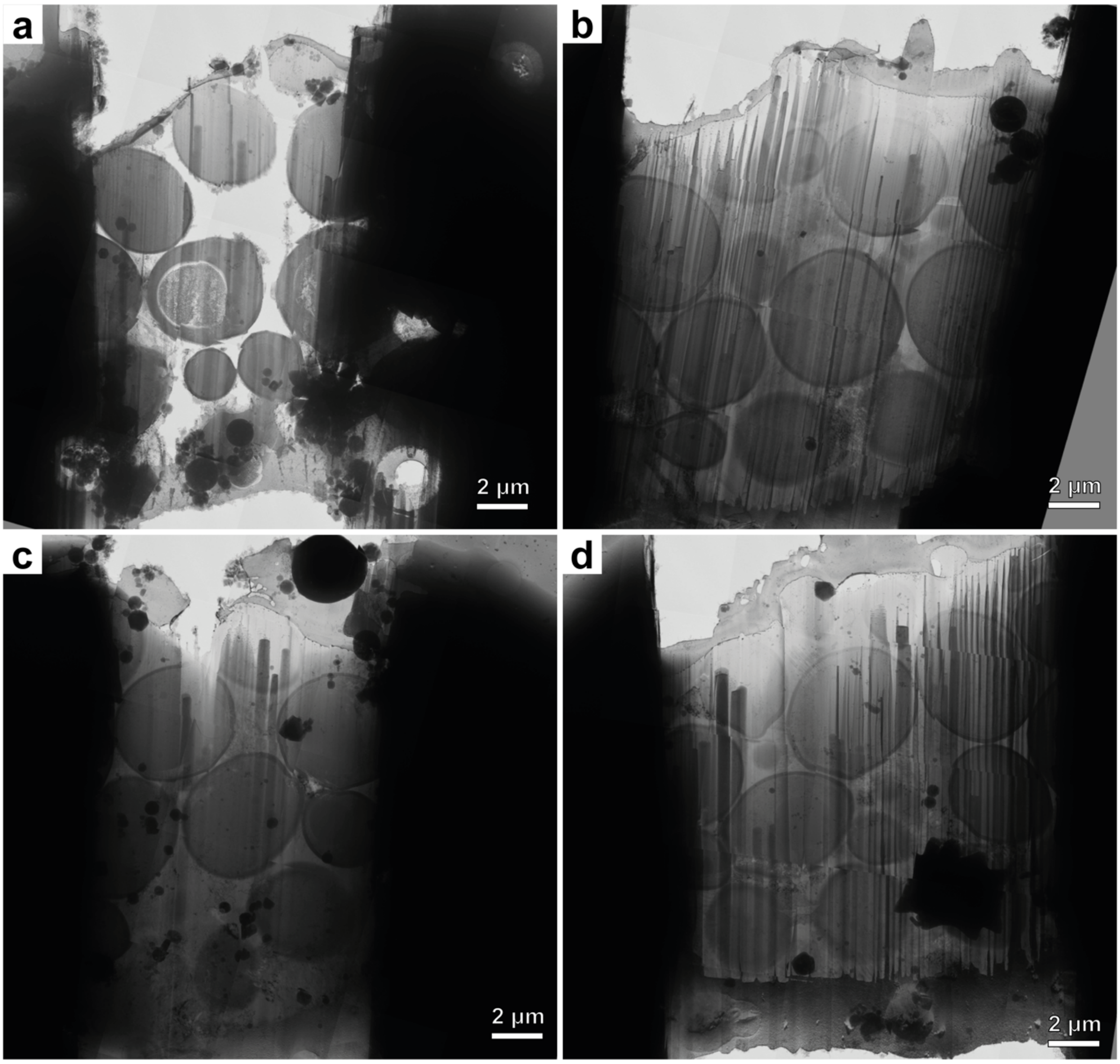


**Supplementary figure 2: TEM overviews of lamellae prepared using ion knife probe exposures. a-d,** Several example overviews of lamellae milled with multiple exposure spots of an ion knife as described in Figure 5, demonstrating the increased curtaining faced with the current implementation of ion knife milling.

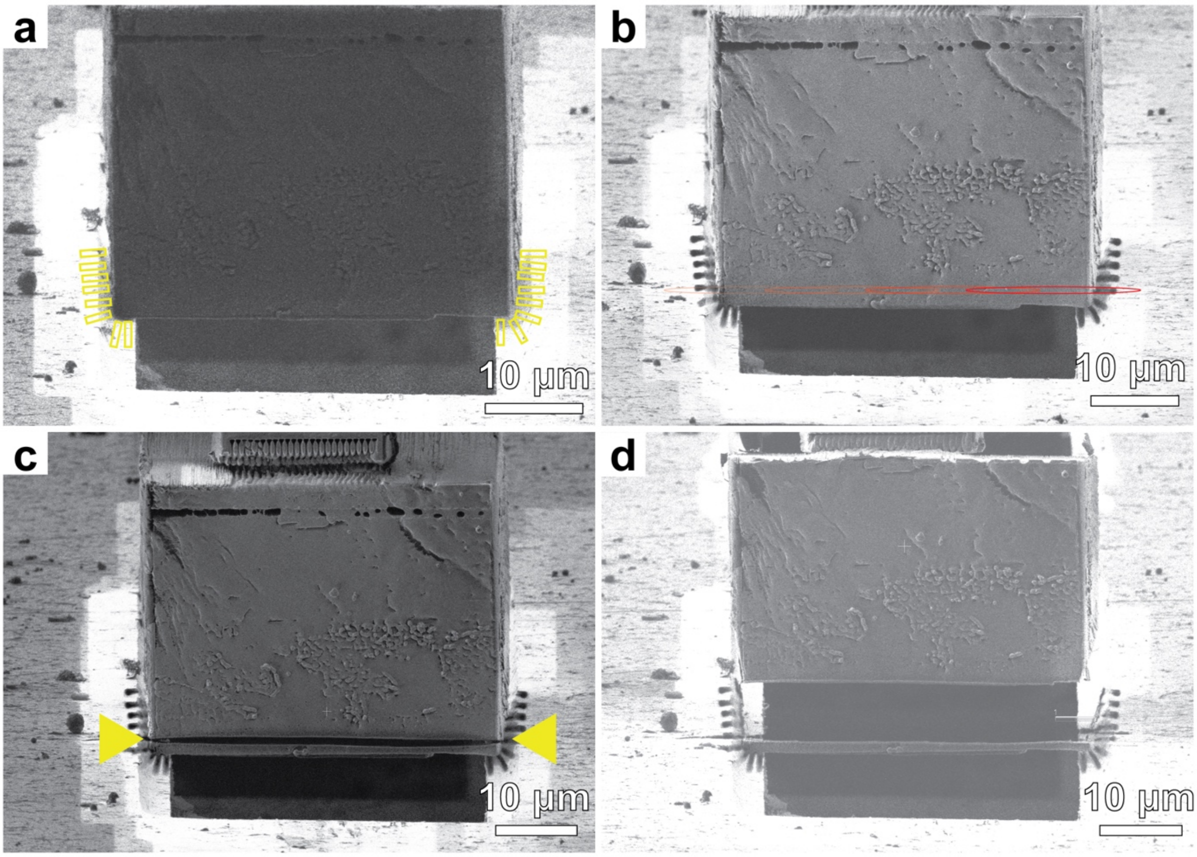


**Supplementary figure 3: Sectioning of lamella in a Serial Lift-Out experiment. a,** Lift-out block aligned with the receiver grid before attachment. Yellow rectangles indicate single-pass regular cross-section patterns that were used for attachment via redeposition. **b,** Image of the block after attachment of the lower end to the receiver grid. Red ovals indicate the ion knife shape and exposure positions. **c,** Image after sectioning using ion knife exposures. Yellow arrowheads indicate the sectioning cut. **d,** After milling of the side attachments of the sectioned lamella, the lift-out block can be moved upwards, the stage can be adjusted to a new position, and a new attachment and sectioning cycle can be performed.
